# Gut microbiota composition in colorectal cancer patients is genetically regulated

**DOI:** 10.1101/2022.03.16.484560

**Authors:** Francesca Colombo, Oscar Illescas Pomposo, Sara Noci, Francesca Minnai, Giulia Pintarelli, Angela Pettinicchio, Alberto Vannelli, Luca Sorrentino, Luigi Battaglia, Maurizio Cosimelli, Tommaso A. Dragani, Manuela Gariboldi

## Abstract

The risk of colorectal cancer (CRC) depends on environmental and genetic factors. Among environmental factors, an imbalance in the gut microbiota can increase CRC risk. Also, microbiota is influenced by host genetics. However, it is not known if germline variants influence CRC development by modulating microbiota composition. We investigated germline variants associated with the abundance of bacterial populations in the normal (non-involved) colorectal mucosa of 93 CRC patients and evaluated their possible role in disease. Using a multivariable linear regression, we assessed the association between germline variants identified by genome wide genotyping and bacteria abundances determined by 16S rRNA gene sequencing.

We identified 37 germline variants associated with the abundance of the genera *Bacteroides, Ruminococcus, Akkermansia, Faecalibacterium* and *Gemmiger* and with alpha diversity. These variants are correlated with the expression of 58 genes involved in inflammatory responses, cell adhesion, apoptosis and barrier integrity. Genes and bacteria appear to be involved in the same processes. In fact, expression of the pro-inflammatory genes *GAL, GSDMD* and *LY6H* was correlated with the abundance of *Bacteroides*, which has pro-inflammatory properties; abundance of the anti-inflammatory genus *Faecalibacterium* correlated with expression of KAZN, with barrier-enhancing functions.

Both the microbiota composition and local inflammation are regulated, at least partially, by the same germline variants. These variants may regulate the microenvironment in which bacteria grow and predispose to the development of cancer. Identification of these variants is the first step to identifying higher-risk individuals and proposing tailored preventive treatments that increase beneficial bacterial populations.

**Authors summary:** Genetic variants describe the variation in the DNA sequence in our genomes and are unique for each person. These variants modify the risk of developing colorectal cancer (CRC) by regulating genes that participate in CRC-associated mechanisms. CRC risk is also affected by microbiota (the microorganisms residing in ourselves). A balanced microbiota helps perform our normal body functions, but can induce cancer, if this balance is lost. Microbiota is affected by factors such as pollution and diet, but is also regulated by genetic variants. However, can genetic variants predispose to cancer risk by regulating microbiota? To answer this question, we sequenced the genetic variants of 93 CRC patients and examined the composition of their intestinal microbiota. We identified variants that regulate the presence of benefic or pathogenic bacteria. The same variants also affect the expression of genes that participate in inflammation, immunity and integrity of intestinal tissue. We found that genetic variants regulate gene expression and microbiota at the same time, predisposing to a higher or lower CRC risk. People with variants predisposing to a higher risk may be benefitted by tailored preventive treatments that increase beneficial bacteria.

## Introduction

Colorectal cancer (CRC) is the fourth most diagnosed tumor and the second leading cause of cancer-related deaths in the world [1]. Its incidence is increasing, especially in developing countries that are undergoing modifications in lifestyle [2]. The vast majority of CRCs are considered to be sporadic [3]. Numerous studies indicated that sporadic CRC is the result of a complex interplay of genetic variants and environmental factors (reviewed in) [4]. Genome-wide association studies have provided evidence for 53 unique CRC susceptibility loci across ethnicities [5]. Among environmental factors, the gut microbiota has emerged as important for some cancers, including CRC, where an imbalance in its composition can contribute to the development of the disease [6]. Interacting closely with host epithelial cells, the microbiota can influence colorectal carcinogenesis via a variety of mechanisms, including microbe-derived factors.

Human gut microbiota composition and host metabolic functions are intimately related and mutually regulated [7]. Studies that investigated whether the microbiota composition is influenced by host genetics highlighted some degree of heritability [8, 9]. Genome-wide association studies on host genetic factors associated with the microbiome composition have led to the identification of several microbial quantitative trait loci (mbQTLs). These mbQTLs are often associated with host genes that participate in nutrition-related metabolic pathways and in immune traits such as barrier integrity or inflammation [8, 10]. Moreover, a study of inflammatory bowel disease (IBD) patients suggested that genetic variants associated with microbiota dysregulation can also affect the immune system and induce inflammation [11]. These observations are of great importance in colorectal cancer, where both inflammation and certain bacterial populations play a role in tumorigenesis [12, 13]. However, no studies associating bacterial populations with genomic variants in CRC patients have been done to date.

We characterized the microbiota in normal colonic mucosa from CRC patients, to identify bacterial populations regulated by host germline variants. Since the vast majority of genetic variants mapped outside coding regions, we looked for possible regulatory effects (i.e. affecting gene expression or splicing) of the identified mbQTLs, with the aim of understanding their functional role.

## Results

Genomic DNA was obtained from resected colorectal mucosa from 95 patients with CRC and subjected to microbiota profiling and SNP genotyping. Genotyping failed in one case and another sample was excluded due to a low call rate. Therefore, mbQTL analyses were done for 93 patients (**Table 1)**. The patients had a median age of 64 years, 60% were men, and 50% had never smoked. The tumor affected the rectum in 59% of cases. There was a broad distribution of pathological stage, with a mode of III (35% of cases). Finally, most patients (85%) had not received neo-adjuvant treatment.

**Table 1.** Clinical characteristics of 93 patients with colorectal cancer.

| Characteristic | Value |
| --- | --- |
| Age at surgery, median (range), years | 64 (38-86) |
| Sex, n (%) |  |
| *Male* | 56 (60) |
| *Female* | 37 (40) |
| Smoking habit, n (%) |  |
| *Ever smoker* | 38 (41) |
| *Never smoker* | 47 (50) |
| *Not available* | 8 (9) |
| Tumor site, n (%) |  |
| *Colon* | 38 (41) |
| *Rectum* | 55 (59) |
| Pathological stage, n (%) |  |
| *I* | 23 (25) |
| *II* | 23 (25) |
| *III* | 33 (35) |
| *IV* | 11 (12) |
| *Not available* | 3 (3) |
| Neo-adjuvant therapy, n (%) |  |
| *Yes* | 14 (15) |
| *No* | 79 (85) |

### Microbiota profiling of non-involved colorectal mucosa

Bacteria associated with patients’ non-involved colorectal mucosa were identified by 16S rRNA gene sequencing, and species diversity was expressed with alpha and beta metrics. The mean Shannon index was higher in V1-V2-V3 16S rRNA gene sequencing data than in V4-V5-V6 data (mean, 6.2 vs. 5.6; *P* = 0.001) (**S1 Table**). In contrast, the mean number of observed OTUs and mean Chao1 estimator were significantly lower in V1-V2-V3. Using Bray-Curtis dissimilarity to estimate beta diversity, we detected significant differences between V1-V2-V3 and V4-V5-V6 data with ANOSIM (*P* = 0.001) and ADONIS (*P* = 0.001). Considering the differences between V1-V2-V3 and V4-V5-V6 datasets in both alpha and beta diversities, we decided to analyze the V1-V2-V3 and V4-V5-V6 datasets separately in this study.

Alpha diversity comparisons between patients grouped according to the clinical characteristics age at surgery, sex or smoking habit showed no differences using either the V1-V2-V3 or V4-V5-V6 dataset. Differently, when patients were grouped according to tumor site, the median Chao1 estimator in the V4-V5-V6 dataset was lower for patients with tumors in the colon than in the rectum (183 vs. 194, respectively, *P* = 0.049; **Fig 1A**). Similarly, the median number of observed OTUs was lower in patients with colonic tumors than with rectal tumors (181 vs. 193, respectively, *P* = 0.049; **Fig 1B**).

**Fig 1.**
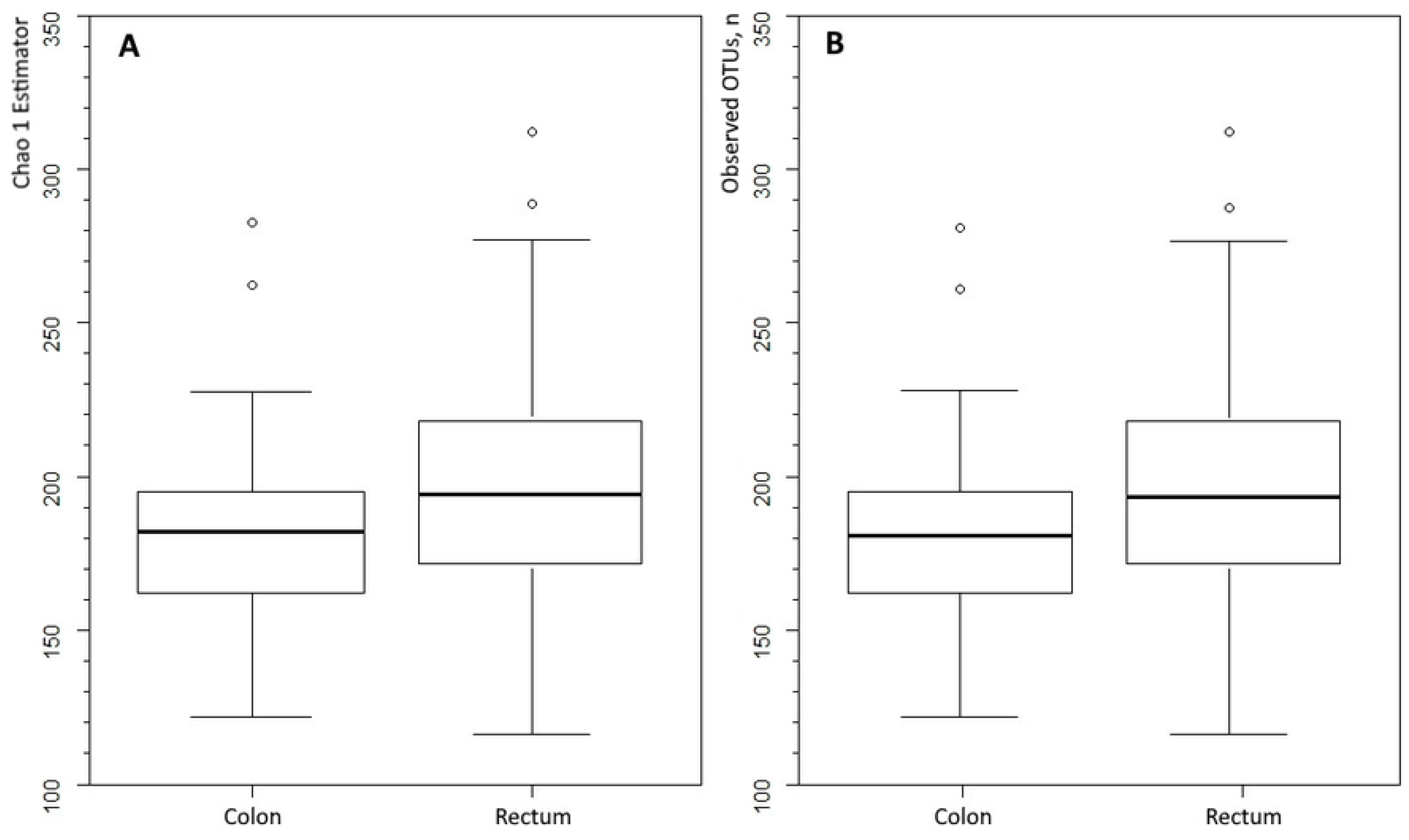
Microbiota diversity in resected colorectal mucosa from patients with colorectal cancer. Data grouped by tumor site, according to the V4-V5-V6 dataset. (**A**) Chao1 estimator, *P* = 0.049, Kruskal-Wallis rank-sum test with continuity correction. (**B**) Number of observed operational taxonomic units (OTUs), *P* = 0.049. The box marks the interquartile range, and the central horizontal line marks the median. Outliers (extreme values, >1.5 times the interquartile range) are shown as circles.

A total of 469 OTUs at genus level was found in both V1-V2-V3 and V4-V5-V6. After data filtering, there were 13 and 12 OTUs, respectively, for analysis. Eight OTUs (i.e. *Bacteroides, Blautia, Coprococcus, Dorea, Faecalibacterium, Pseudomonas, Roseburia*, and *Ruminococcus* genera) were identified in both datasets, for 17 unique OTUs (**Table 2**). In multivariate linear regression, seven unique OTUs independently associated with patients’ characteristics, namely age at surgery (three OTUs), smoking habit (three OTUs), and tumor site (two OTUs). The relative abundance of *Roseburia* associated with age and smoking habit, with lower abundance in older patients (beta, −0.050 and −0.045 for V1-V2-V3 and V4-V5-V6 datasets, respectively) and greater abundant in ever smokers (beta = 0.95). *Colinsella* and *Parabacteroides* OTUs were also more abundant in ever smokers than never smokers (beta, 1.1 and 0.89, respectively). The abundance of *Dorea* decreased with age, whereas that of *Escherichia* increased. Finally, *Pseudomonas* was more abundant and *Gemmiger* less abundant in patients who had a rectal than colonic tumor.

**Table 2.** Association of operational taxonomic units (OTUs) and clinical characteristics. OTUs observed in >25% of samples and with mean relative abundance >1%, and associations between the clr-transformed relative abundance and patient characteristics, by multivariate linear regression

| OTU | V1-V2-V3<br>Dataset <sup>a</sup> | V4-V5-V6<br>Dataset | Clinical<br>characteristic | Beta (*P*)<sup>b</sup> |
| --- | --- | --- | --- | --- |
| *Akkermansia* | N | Y | None | - |
| *Alistipes* | Y | N | None | - |
| *Bacteroides* | Y | Y | None | - |
| *Blautia* | Y | Y | None | - |
| *Collinsella* | N | Y | Smoking | 1.1 (0.045)<sup>2</sup> |
| *Coprococcus* | Y | Y | None | - |
| *Dorea* | Y | Y | Age at surgery | -0.046 (0.023)<sup>2</sup> |
| *Escherichia* | Y | N | Age at surgery | 0.079 (0.00074)<sup>1</sup> |
| *Faecalibacterium* | Y | Y | None | - |
| *Gemmiger* | N | Y | Tumor site<sup>c</sup> | -0.97 (0.033)<sup>2</sup> |
| *Parabacteroides* | Y | N | Smoking | 0.89 (0.029)<sup>1</sup> |
| *Prevotella* | Y | N | None | - |
| *Pseudomonas* | Y | Y | Tumor site<sup>c</sup> | 0.93 (0.030)<sup>2</sup> |
| *Robinsoniella* | N | Y | None | - |
| *Roseburia* | Y | Y | Smoking | 0.95 (0.035)<sup>1</sup> |
|  |  |  | Age at surgery | -0.050 (0.0063)<sup>1</sup> |
|  |  |  | Age at surgery | -0.045 (0.014)<sup>2</sup> |
| *Ruminococcus* | Y | Y | None | - |
| *Stenotrophomonas* | Y | N | None | - |

<sup>a</sup> Sequence data from the six hypervariable regions were aggregated into two datasets (V1-V2-V3 and V4-V5-V6) and analyzed separately. <sup>b</sup> Beta and *P*-value from <sup>1</sup>V1-V2-V3 or <sup>2</sup>V4-V5-V6 datasets. <sup>c</sup> Rectum/colon

We also observed that many OTU abundances were significantly correlated among each other (Pearson’s |r| > 0.20 and *P* <0.05; **S1 Fig**). In the V1-V2-V3 dataset, the number of correlations per OTU ranged from three for *Blautia* to 11 for *Faecalibacterium, Alistipes*, and *Dorea*. In the V4-V5-V6 dataset, the number of correlations per OTU ranged from three for *Blautia* to 10 for *Roseburia, Ruminococcus*, and *Faecalibacterium*. In both datasets, more correlations were positive than negative.

### Association of germline variants with microbiota-related quantitative traits

Genome-wide genotyping on Axiom Precision Medicine Research Arrays provided data on 856,427 variants. Of these, 17,919 were removed since they had a call rate <98% and 536,891 variants with a MAF <5% were also removed. Therefore, data on 301,884 germline variants were available for analysis.

Multivariable linear regression was used to combine genotype data and microbiota-related quantitative traits to identify mbQTLs. Analysis using the Shannon index identified 12 loci comprising 18 variants (FDR <0.1) on seven unique chromosomes. Of the 18 variants, 10 (on four chromosomes) derived from V1-V2-V3 sequencing data (**Fig 2a; S2 Table**); these variants all had an FDR <0.05, and two of them (rs78578513 and rs5994535 on 22q12.3) almost reached the genome-wide significance threshold of nominal *P* = 5.0 x 10^-8^. The remaining eight variants derived from V4-V5-V6 sequencing (**Fig 2b; S2 Table**); these variants were distributed on seven chromosomes and had an FDR <0.1. For all these mbQTLs, there was a negative correlation between the number of minor alleles in the genotype and the Shannon index, indicating a reduction in microbial diversity with an increasing number of minor alleles. No mbQTLs were found when the Chao1 estimator and number of observed OTUs were analyzed.

**Fig 2.**
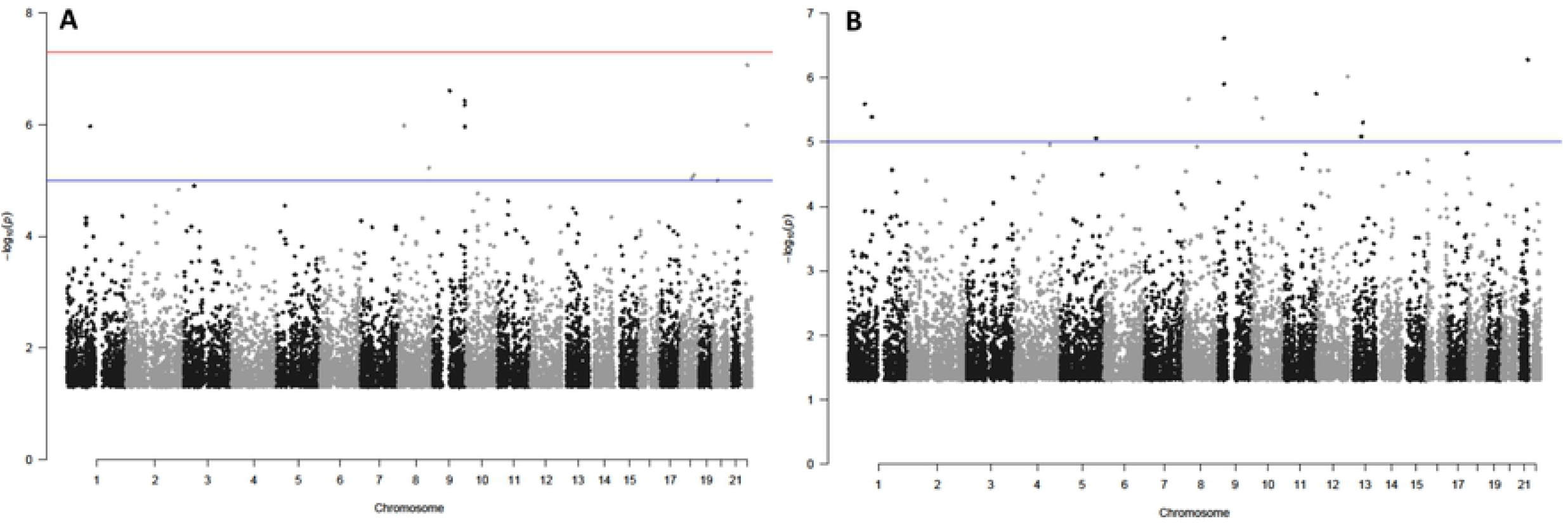
Manhattan plots of Shannon index-specific mbQTLs. (**A**) V1-V2-V3 dataset. (**B**) V4-V5-V6 dataset. Red horizontal line, threshold for genome-wide significance (*P* = 5.0×10^-8^). Blue horizontal line, FDR=0.10.

When multivariable linear regression was run using the clr-transformed relative abundance of OTUs as the microbiota-related quantitative trait, we identified 37 unique mbQTLs in 23 loci on 11 chromosomes (FDR <0.10) (**S2 Table**). *Bacteroides* had the highest number of unique mbQTLs (n = 28), including four (rs7815797, rs72691617, rs7817574, rs79744701) at 8q24.3 that were found in both V1-V2-V3 and V4-V5-V6 datasets (**Fig 3A, B**). A single mbQTL was associated with *Ruminococcus* abundance in the V1-V2-V3 dataset (**Fig 3C**). Another eight mbQTLs in seven loci were identified as being associated with *Akkermansia, Faecalibacterium* or *Gemmiger* abundance (**Fig 3D-F**). The most significant association overall (*P* = 1.23 x 10^-9^) was observed between *Akkermansia* and rs4527077 on chromosome 8q24.22. This variant, together with its neighbor rs4736470 and with rs12472313 on chromosome 2, were the only three polymorphisms that correlated directly with microbial abundance, meaning that an increasing number of minor alleles of these single nucleotide polymorphisms (SNPs) associated with more abundant *Akkermansia*. All the other mbQTL variants correlated inversely with microbial abundance. Among the mbQTLs found, only two have so far been reported to be associated with a specific trait, according to the NHGRI-EBI Catalog of Human Genome-Wide Association Studies. In particular, variants rs79744701 (G>A) and rs7817574 (T>C) were both found to associate with serum levels of apolipoprotein A1 and HDL cholesterol [14, 15].

**Fig 3.**
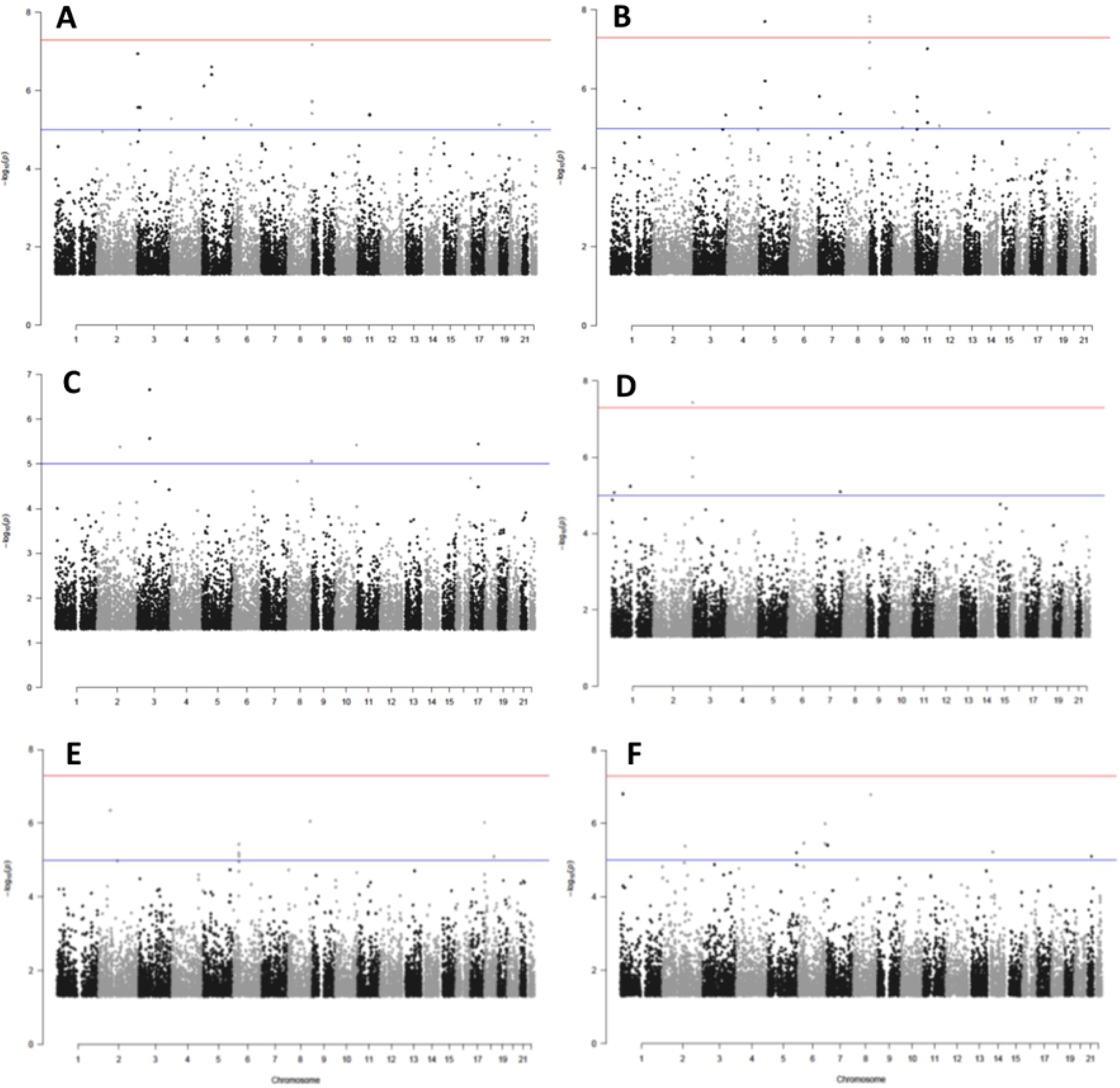
Manhattan plots of OTU-specific mbQTLs. (**A**) *Bacteroides* V1-V2-V3, (**B**) *Bacteroides* V4-V5-V6, (**C**) *Ruminococcus* V4-V5-V6, (**D**) *Akkermansia*, V4-V5-V6 (**E**) *Faecalibacterium* and (**F**) *Gemmiger* V4-V5-V6. Red horizontal line, threshold for genome-wide significance (*P* = 5.0×10^-8^). Blue horizontal line, FDR=0.10

### Regulatory effects of mbQTLs in non-involved colorectal mucosa

Most identified mbQTLs mapped outside coding regions. To determine if the mbQTLs have genetic regulatory effects, we interrogated GTEx and Ensembl VEP databases and identified 58 genes whose expression or splicing (in any tissue) had been reported to be associated with the variants we identified as colorectal mbQTLs. With our Shannon index-specific mbQTLs, we identified 28 genes whose expression (n = 24) or splicing (n = 9) varied according to genotype (**S3 Table**). In particular, two genes were upregulated and two downregulated in colorectal mucosa.

With our OTU-specific mbQTLs, we identified 30 genes regulated by genotype (**Table 3**). We found 25 genes linked to *Bacteroides*-specific mbQTLs, including one upregulated and two downregulated in colorectal mucosa. We also found three genes with *Akkermansia-*specific and one each with *Faecalibacterium-* and *Gemmiger*-specific mbQTLs. Of the 58 identified genes, 29 are known to be differentially expressed in colorectal adenocarcinoma, with respect to normal tissue, according to The Cancer Genome Atlas database (**S4 Table**). Of these, 10 were associated with the Shannon index and 19 associated with *Bacteroides, Akkermansia* or *Gemmiger* abundance in this study.

**Table 3.** Genetic regulatory effects of operational taxonomic unit (OTU)-specific mbQTLs.

| OTU | SNPs | Gene | Locus | Expression change<sup>1</sup> | Splicing change |
| --- | --- | --- | --- | --- | --- |
| Bacteroides | rs111574827, rs3790893 | ADGRL2 | 1p31.1 | - | IR |
|  | rs12563507 | ATP1B1 | 1q24.2 | ↑ | - |
|  | rs12563507 | NME7 | 1q24.2 | ↑ | AS |
|  | rs12563507 | CCDC181 | 1q24.2 | - | AS |
|  | rs711582 | LRRN1 | 3p26.1 | ↑* | IR |
|  | rs34222640 | ITPR1 | 3p26.1 | - | IR |
|  | rs78482149 | TP63 | 3q28 | - | IR |
|  | rs78482149, rs56201661 | IL1RAP | 3q28 | ↑ | - |
|  | rs28368180, rs35361432, rs116871956 | SLC38A9 | 5q11.2 | ↓* | AS, IR |
|  | rs75284212 | RP11-274B21.2 | 7q32.1 | ↓ | - |
|  | rs72691617, rs7817574, rs79744701 | LY6H | 8q24.3 | ↓ | - |
|  | rs7815797, rs72691617, rs7817574, rs79744701 | GPIHBP1 | 8q24.3 | ↓* | - |
|  | rs79744701 | ZFP41 | 8q24.3 | ↓ | - |
|  | rs79744701 | GLI4 | 8q24.3 | ↑ | - |
|  | rs79744701 | MINCR | 8q24.3 | ↑ | - |
|  | rs79744701 | TOP1MT | 8q24.3 | ↑ | AS |
|  | rs7815797, rs72691617, rs7817574 | ZC3H3 | 8q24.3 | ↓ | - |
|  | rs7815797, rs72691617, rs7817574 | GSDMD | 8q24.3 | ↓ | - |
|  | rs7815797, rs72691617, rs7817574 | RP11-661A12.5 | 8q24.3 | ↓ | - |
|  | rs3833782 | RPL27A | 11p15.4 | - | IR |
|  | rs3833782, rs1104774 | STK33 | 11p15.4 | ↓ | AS |
|  | rs3833782, rs1104774 | DENND2B | 11p15.4 | - | IR |
|  | rs74830761 | GAL | 11q13.3 | ↓ | - |
|  | rs4901170 | RP11-280K24.4 | 14q22.1 | - | AS |
|  | rs4901170 | GNG2 | 14q22.1 | - | IR |
| Akkermansia | rs598300 | ROCK1P1 | 18p11.32 | ↑ | - |
|  | rs598300 | USP14 | 18p11.32 | ↓ | - |
|  | rs598300 | THOC1 | 18p11.32 | - | IR |
| Faecalibacterium | rs7526230 | KAZN | 1p36.21 | - | IR |
| Gemmiger | rs73129818 | IQCA1 | 2q37.3 | ↓↑ | - |

<sup>1</sup> Upregulation (↑) or downregulation (↓) in the colon (\*) or other tissues, associated with an increasing frequency of minor alleles (GTEx database; FDR<0.05). ↓↑ Divergent results in
two tissues; -, no change reported. AS, alternative splicing-derived isoforms (GTEx database); IR, intron retention (Ensembl VEP)

Ontology annotations in the DAVID Bioinformatic Database indicate that the products of 23 of the 58 genes localize to the membrane, 10 are extracellular, and 17 are glycosylated (**S5 Table)**. Also, four genes (*GAL, GSDMD, IL1RAP* and *ITGB2*) are associated with the inflammatory response, three with cell adhesion (*ATP1B1, LOXL2* and *ITGB2*) and three with apoptosis (*THOC1, TP63* and *ITGB2*) **(S6 Table**). Four variants associated with the Shannon index and seven with *Bacteroides*, as well as one variant for *Akkermansia, Faecalibacterium* and *Gemmiger* each, map in predicted promoter or enhancer regions or in CTCF (CCCTC-binding factor) binding sites (**S7 Table**). The single variant associated with *Ruminococcus* has no reported effects on the expression of any gene nor is found in a regulatory region. Finally, we observed the pathogenic species *Bacteroides fragilis* in 36 patients, with an abundance greater than 1% in 13 of them, according to data from 16S rRNA gene V1-V2-V3 regions (**S8 Table**).

## Discussion

This study investigated associations between the microbiota of non-involved colorectal mucosa in CRC patients and both patients’ clinical characteristics and genetic variants. We found that changes in abundance of the genera *Dorea, Roseburia* and *Escherichia* were associated with age at surgery, *Gemmiger* with tumor site, and *Collinsella* and *Parabacteroides* with smoking habit. Moreover, changes in the abundance of *Bacteroides, Akkemansia, Faecalibacterium* and *Gemmiger* correlated with germline variants already known to be associated with the expression of 30 genes, while germline variants associated with the Shannon index were correlated with 28 genes. These genes participate in mechanisms of great relevance in cancer development, namely the inflammatory response, apoptosis, cell adhesion and epithelial barrier function, and 29 are known to be dysregulated in CRC. Interestingly, similar mechanisms were associated with *Bacteroides, Akkermansia* or *Gemmiger* abundance.

The risk and incidence of CRC and inflammatory diseases are higher in older patients [1]. Accordingly, we found that two genera with anti-inflammatory properties, namely the short-chain fatty acid producers *Roseburia* and *Dorea* [16, 17], were negatively associated with age, while the potentially pro-inflammatory *Escherichia* was positively associated [18, 19]. A smoking habit was positively associated with the abundance of *Collinsella, Parabacteroides* and *Roseburia* genera. *Collinsella* has been reported to increase mucosal barrier permeability and induce the expression of pro-inflammatory IL-17 and NFkB1 in patients with rheumatoid arthritis [20]. Comparing patients with rectal vs. colonic tumors, we found that the opportunistic pathogen *Pseudomonas* [21], was more abundant in the rectum. Instead, *Gemmiger* was more abundant in the colon; the abundance of this genus has been shown to be reduced in patients with inflammatory diseases [22]. These observations are consistent with findings that the gut bacterial composition varies along the gastrointestinal tract [23].

This study found 18 mbQTLs associated with bacterial diversity (Shannon index) and 37 mbQTLs associated with the abundance of *Bacteroides, Akkemansia, Faecalibacterium, Gemmiger* and *Ruminococcus*. Genes associated with mbQTLs were already known to be involved in host metabolism, immunity and cell and tissue barrier integrity [8, 10]. Of all the mbQTLs identified here, 34 were already known to be expression or splicing QTLs (here, referred to as “mbesQTLs”) for 58 genes. The products of 23 of these genes localize to the cell membrane and may alter gut barrier–microbiota interactions. Eleven of these genes are known to participate in immune mechanisms, including the innate response, inflammation and barrier integrity [24]. Additionally, six genes are known to be involved in apoptosis and transcription regulation, two pathways commonly dysregulated in cancer.

Our study suggests that a pro-inflammatory environment reduces diversity of the normal colonic mucosa-associated microflora, as already reported for IBD [25]. Indeed, among the 28 genes with Shannon index-associated mbesQTLs, *BPIFC* (bactericidal/permeability-increasing fold-containing family C protein), *PIK3IP1* (phosphoinositide-3-kinase-interacting protein 1), *ITGB2* (integrin subunit beta 2), and *TRPM3* (transient receptor potential cation channel subfamily M member 3) are involved in inflammation and immune mechanisms [26–30]. In particular, *BPIFC* belongs to a family of genes expressing antibacterial peptides that are released by neutrophils and bind lipopolysaccharides [31]. PIK3IP1 is a transmembrane protein that inhibits T cell responses to tumoral cells and intracellular bacteria [32, 33]. The presence of these mbesQTLs corresponds to a lower Shannon index, lower expression of the anti-inflammatory *PIK3IP1* gene and higher expression of the pro-inflammatory *BPIFC* gene. Genes involved in immunity were also identified with the mbesQTLs associated with *Bacteroides, Akkermansia* and *Faecalibacterium* genera. *Bacteroides* comprises pathogenic species with pro-inflammatory and invasive properties; they are able to colonize epithelial cells and their abundance is increased in CRC [6, 34, 35]. *Bacteroides fragilis* has a known role in colon carcinogenesis [36, 37].

We observed germline variants in 5 genes predisposing to a more efficient barrier and lower inflammation that would hinder *Bacteroides* expansion. *GAL* (galanin and GMAP prepropeptide) encodes for two proteins: galanin and galanin message-associated peptide (GMAP). Galanin regulates intestinal motility and induces chloride ion secretion in pathogen-derived inflammation-associated diarrhea [38], while GMAP expression increases in response to lipopolysaccharide (an inducer of inflammation) [39]. Gasdermin D (*GSDMD*) is the main catalyst of pyroptosis, a cellular programmed necrosis induced by the presence of intracellular lipopolysaccharide [40]. IL-1 receptor accessory protein (*IL1RAP*) is essential for IL-1 signaling [41]. It is associated with IL-1 receptor (IL-1R1), which participates in CRC-associated inflammation and tumor progression [42]. *LY6H* (lymphocyte antigen 6 family member H) is a pro-inflammatory regulator that inhibits the nicotinic acetylcholine receptor [43], which is a negative regulator of innate immunity antimicrobial peptides (AMPs) [44]. AMPs are potent pro-inflammatory molecules with a broad antibacterial spectrum, central to the innate epithelial immune response, and their expression can be regulated by the microbiota itself [45, 46]. The product of *ATP1B1* (ATPase Na+/K+ transporting subunit beta 1) upregulates the expression of tight-junction proteins and increases the epithelial barrier function in the lung [47]. The abundance of *Bacteroides* was lower in the presence of all its associated mbesQTLs. The expression of the pro-inflammatory genes *GAL, GSDMD* and *LY6H* was also reduced. *ATP1B1* and *IL1RAP* were instead more expressed.

*Akkermansia muciniphila*, the most abundant *Akkermansia* species in the human gut, is a mucin degrader considered beneficial for its anti-inflammatory and gut barrier function-enhancing properties [48–50]. However, it is enriched in CRC [51]. We observed an increase in *Akkermansia* abundance and *USP14* (ubiquitin-specific protease 14) expression in the presence of the mbesQTL. *USP14* is often overexpressed in cancer [52], and its product activates NF-κB and ERK1/2 in response to microbial infection [53]. Interestingly, NF-κB and ERK1/2 induce mucin expression [54], which may explain the observed increase in *Akkermansia* abundance.

*Faecalibacterium prausnitzii*, the *Faecalibacterium* species most represented in the human gut, has anti-inflammatory properties and is reduced in various intestinal disorders [55, 56]. Kazrin (*KAZN*), which acts as a cytolinker by functionally connecting adherens junctions and desmosomes [57], was identified with the mbesQTL associated with *Faecalibacterium* and presents a splicing variation with unknown function.

A search of the 58 genes within The Cancer Genome Atlas revealed that 10 of the 28 genes identified with Shannon index-associated mbesQTLs and 19 of the 29 mbesQTLs associated with *Bacteroides, Akkermansia* and *Gemmiger* are dysregulated in CRC. Among the genes associated with mbesQTLs and known to be differently expressed in colorectal adenocarcinoma tissue according to The Cancer Genome Atlas database, *USP14, LOXL2* (lysyl oxidase-like 2) and *TP63* (tumor protein 63) link the microbiota composition with cancer development. *LOXL2* is involved in extracellular matrix (ECM) remodeling [58], and is predicted to have intron retention with unknown function in presence of the Shannon index-associated mbesQTL. Changes in ECM structure or composition can promote inflammation due to bacterial invasion that can lead to tumor development [59]. *TP63* expresses a tumor suppressor from the p53 family and an important regulator of cell differentiation, proliferation and survival [60]. It also enhances CXCR4 expression [61], which can be activated by various pathogens such as *Porphyromonas gingivalis* and *Chlamydia pneumoniae* to modulate the immune response and facilitate infection [62, 63]. We found an intron retention with unknown function in *TP63* associated with *Bacteroides* mbQTLs.

We observed the presence of the *B. fragilis* in 36 patients. While other *Bacteroides* species are commensals normally present in healthy microbiota, *B. fragilis* is a pathogen. It is the most common anaerobe isolated from extraintestinal infections, and it has pro-inflammatory activities and contributes to colon carcinogenesis [36]. However, this OTU was not associated with clinical data in this study.

In conclusion, we identified mbesQTLs associated with genes involved in the regulation of the inflammatory environment and with the abundance of bacterial populations with immune-modulatory properties. We observed a correlation between pro-inflammatory genes and the abundance of the pro-inflammatory *Bacteroides*, while an inverse correlation was observed for barrier-reinforcing genes. mbesQTLs also showed a correlation between the mucin-degrading *Akkermansia* and expression of genes that regulate mucin expression. mbesQTL-associated genes may regulate the microenvironment in which bacteria grow and predispose to the development of cancer, some of these genes are also known to participate in carcinogenesis. Identifying variants that promote the growth of disease-associated bacterial populations may help find higher-risk individuals who could benefit from preventive treatment such as antibiotics that inhibit cancer-associated bacteria and supplementation with commensal bacteria (probiotics) or metabolites generated by microbiota (postbiotics). Further studies with larger sample sizes and control populations are needed to clarify whether genetic variants associated with the regulation of both intestinal microbiota composition and gene expression have a role in CRC development.

## Materials and methods

### Ethics statement

This case series is part of a broader research program on colorectal cancer, for which a biobank of lung tissue and extracted nucleic acids is maintained at Fondazione IRCCS Istituto Nazionale dei Tumori, Milan. The protocol for collecting tissue specimens and related clinical data was approved by the ethics committee of the Fondazione IRCCS Istituto Nazionale dei Tumori, Milan. All patients who donated samples provided written informed consent for the use of their materials for research purposes.

### Patient series and biological material

The study included 95 patients with CRC who underwent surgery at the Colorectal Surgery Unit of Fondazione IRCCS Istituto Nazionale dei Tumori between 2009 and 2010 in the context of a larger study on CRC genetics [64].

Patients with CRC at any pathological stage, determined following the American Joint Committee on Cancer (AJCC) TNM system [65], were eligible for inclusion in the study. To avoid potential consequences of inflammation on gut microbiota, patients were excluded from this study if signs of acute or chronic colorectal inflammation or stenosis were observed during surgery. Furthermore, patients were excluded if they had received antibiotics in the 6 months before surgery. Clinical data were collected regarding age at surgery, sex, self-reported smoking habit (scored as ever or never smoker of tobacco-containing products), tumor site (colon or rectum), pathological stage and whether the patients received neo-adjuvant therapy before surgery (yes/no).

As per standard operating procedures, patients underwent bowel preparation with 2 sachets of sodium picosulfate (4 hours apart in the evening before surgery) and received cefazolin (2 g intravenously) and metronidazole (500 mg intravenously) 40 minutes before incision. At the end of surgery, biopsies were collected from non-involved mucosa as far as possible from the tumor in the resected colorectal tissue. The samples were placed in RNAlater Stabilization Solution (Sigma-Aldrich) and then used for DNA extraction with the DNeasy Blood & Tissue Kit (Qiagen). Genomic DNA was quantified using the NanoDrop 2000c UV–Vis spectrophotometer (Thermo Fisher Scientific), diluted in water and stored at −20 °C until use in microbiota profiling and SNP genotyping.

### Microbiota profiling

Genomic DNA samples from non-involved colorectal mucosa of the 95 patients were used for microbiota profiling by sequencing the bacterial 16S rRNA gene. To amplify the first three hypervariable regions (V1, V2 and V3) of the 16S rRNA gene, we used AmpliTaq Gold DNA Polymerase (Thermo Fisher Scientific) and the 27F and 519R universal primers **(Table 4**) [66, 67]. For the V4, V5 and V6 hypervariable regions, we ran PCR with the 533F and 1100R universal primers [67, 68]. The primers sequences were modified to include Illumina Nextera index PCR adaptor sequences (Illumina).

**Table 4.**
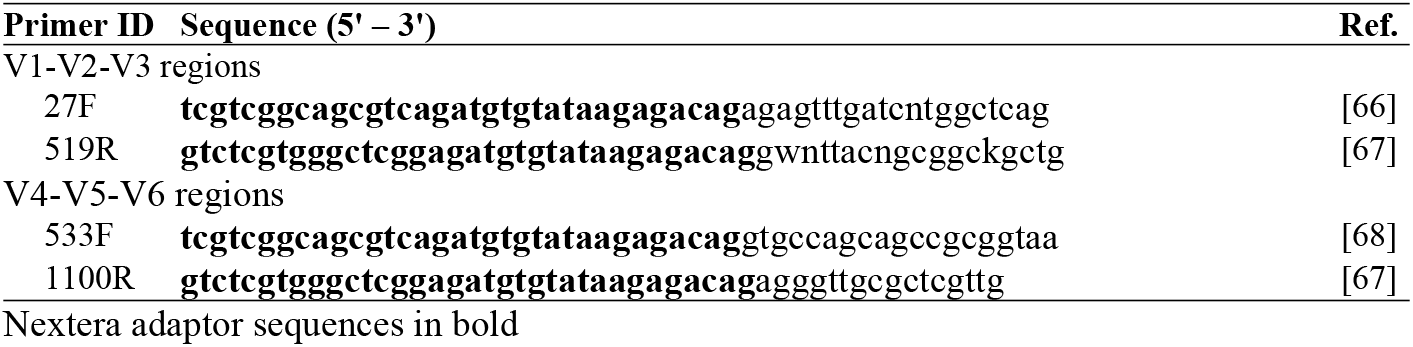
Universal primers used for 16S rRNA amplification.

PCR products were purified and sent to Eurofins Genomics (Edersberg, Germany) for index PCR, pooling and normalization of amplicons, and sequencing on an Illumina MiSeq sequencer with the 2 x 300 bp paired-end read module. Eurofins Genomics also did the read processing (according to primer sequences using proprietary scripts), taxonomy assignation and copy number correction. The company provided the data in files with fasta and biom formats for further analyses in our institutes.

Paired-end read merging was performed using FLASH software [69]. Chimeric reads were identified and removed with UCHIME, implemented with VSEARCH software [70, 71]. Reads were sorted into operational taxonomic units (OTUs) through the minimum entropy decomposition method [72, 73]. Taxonomic assignment was performed with BLAST in QIIME software v. 1.9.1 (http://qiime.org/) [74, 75], using the NCBI_nt database as reference and with a minimum identity of 70% across at least 80% of the OTU representative sequence. OTU abundances were normalized considering their lineage-specific gene copy number [76]. Microbial profiling was carried out separately for V1-V2-V3 and V4-V5-V6 amplicons.

QIIME software was used to analyze bacterial species diversity. Alpha diversity was assessed with the Shannon index, total number of observed OTUs, and the Chao1 estimator [77]. Beta diversity metric was assessed with the Bray-Curtis index of dissimilarity. Analyses of microbiota abundance were performed at the genus and species levels. To analyze OTUs at the genus level, we first removed OTUs missing a genus-level classification. To minimize the number of tests, we only considered taxa present in at least 25% of samples and with mean relative abundance greater than 1% among all samples. To remove compositional constraints typical of microbiota data, zero values were substituted with a pseudo-value less than the lesser relative abundance of the dataset, and relative abundances were transformed using centered log ratio (clr) transformation with the clr function of the Compositions R package [78].

### SNP genotyping

Genomic DNA (100 ng per sample) was subjected to genome-wide genotyping using Axiom Precision Medicine Research Arrays (Thermo Fisher Scientific) on an Affymetrix Gene Titan platform at Eurofins Genomics Europe Genotyping (Galten, DK). Raw data provided by Eurofins were analyzed in our institutes using the Axiom Analysis Suite (Thermo Fisher Scientific). Genotype data were subjected to quality control using PLINK software v1.90b6.16 [79]. Per-sample quality check discarded samples with ≥ 2 % missing genotypes. Genotyped variants with a call rate <98% or a minor allele frequency (MAF) <5% were excluded from analysis.

### Statistical analyses

Alpha diversity metrics were compared between samples grouped by hypervariable regions V1-V2-V3 and V4-V5-V6 using a non-parametric *t*-test, adjusted for multiple testing by calculating the false discovery rate (FDR) using the Benjamini-Hochberg method. Differences in the Bray-Curtis index of dissimilarity were tested for significance with a permutation test with pseudo-F ratios (ADONIS function) and an analysis of similarities (ANOSIM function). These analyses were done in QIIME software v. 1.9.1, and the results were considered statistically significant when *P* <0.01.

To determine if alpha diversity indexes differed between subgroups of patients defined by clinical characteristics (age, sex, smoking habit and tumor site), the non-parametric Kruskal-Wallis rank-sum test with continuity correction was used. To study associations between the clr-transformed relative abundance of OTUs and patient characteristics, we used multivariate linear regression. Pearson’s correlation in relative abundance between OTUs was assessed using the rcorr function of the Hmisc package of R, and correlograms were drawn using R package corrplot. The significance threshold was set at *P* <0.05, and correlation was defined as |r| >0.02.

To identify mbQTLs, we used multivariable linear regression analysis to assess associations between germline polymorphisms and microbiota-related traits, including alpha and beta diversity metrics and clr-transformed relative abundance of OTUs. Sex, age, pathological stage, tumor site, and smoking habit were entered as covariates. The distance to define two loci on the same chromosome as independent loci was set to >1Mb. We adjusted for multiple testing using the Benjamini-Hochberg procedure to obtain the FDR [80], and a significance threshold was set at FDR <0.10. We also consider the commonly accepted genome-wide significance threshold, set at the nominal *P* <5.0×10^-8^ [81]. These analyses were done using PLINK software. Manhattan plots were drawn using the manhattan function of the qqman package in R.

To investigate possible regulatory effects of the identified mbQTLs, we consulted the Genotype-Tissue Expression (GTEx) project database v7 [82], and the Ensembl Variant Effect Predictor (VEP) [83], on 19 April 2021. In GTEx, we looked for matches between our mbQTLs and any expression QTL or splicing QTL in any tissue. Genes associated with these expression or splicing QTLs and genes predicted by VEP to retain an intron on their products in the presence of the minor allele of the identified mbQTLs were considered for analysis. Ontology annotation of these genes was obtained from DAVID Bioinformatic Database v6.8 [24], and information on differential expression was obtained from The Cancer Genome Atlas [84].

## Acknowledgements

The authors acknowledge the contributions of Valerie Matarese, PhD, who provided scientific editing.

## Data availability

The genotyping data that support the findings of this study have been deposited in the European Genome-Phenome Archive with the accession code EGASXXXXXXX. The 16S rRNA gene sequencing data that support the findings of this study are available from https://icedrive.net/s/7RhCk77Fw5jAFw87AzNRixBGa8WA.

## Supporting information

**S1 Fig. Correlation matrices for OTU abundance.** A) V1-V2-V3 dataset. B) V4-V5-V6 dataset. Numbers along the matrix indicate the OTU in the list. Pearson’s correlation coefficients are indicated on a dichromatic scale, and the sizes of the circles are proportional to the correlation coefficient.

**S1 Table. Alpha diversity in V1-V2-V3 and V4-V5-V6 regions**

**S2 Table. Associations between germline variants and microbiome-related quantitative traits.**

**S3 Table. Genetic regulatory effects of Shannon index-specific mbQTLs**

**S4 Table. Genes associated with differently expressed eQTLs in colon or rectum adenocarcinoma versus normal tissue.**

**S5 Table. Functional annotation clustering of genes associated with an mbQTL**

**S6 Table. Classification of 58 genes under genetic regulation by variants in our colorectal mbQTLs.**

**S7 Table. mbQTL variants located in gene regulatory elements according to the Ensembl Variant Effect Predictor**

**S8 Table. Abundance and occurrence of Bacteroides species**

